# Currently available intravenous immunoglobulin (Gamunex^®^-C and Flebogamma^®^ DIF) contains antibodies reacting against SARS-CoV-2 antigens

**DOI:** 10.1101/2020.04.07.029017

**Authors:** José-María Díez, Carolina Romero, Rodrigo Gajardo

## Abstract

**Background:** There is a critical need for effective therapies that are immediately available to control the spread of COVID-19 disease. In this study, we assessed currently marketed intravenous immunoglobulin (IVIG) products for antibodies against human common coronaviruses that may cross-react with the SARS-CoV-2 virus.

**Methods:** Gamunex^®^-C and Flebogamma^®^ DIF (Grifols) IVIG were tested against several betacoronaviruses antigens using ELISA techniques: HCoV (undetermined antigen), HCoV-HKU1 (N protein), SARS-CoV (culture lysate), MERS-CoV (N protein; S1 protein/RBD; S protein), and SARS-CoV-2 (S1 protein).

**Results:** Both IVIG products showed consistent reactivity to components of the tested viruses. Positive cross-reactivity was seen in SARS-CoV, MERS-CoV, and SARS-CoV-2. For SARS-CoV-2, positive reactivity was observed at IVIG concentrations ranging from 100 μg/mL with Gamunex-C to 1 mg/mL with Flebogamma 5% DIF.

**Conclusion:** Gamunex-C and Flebogamma DIF IVIG contain antibodies reacting against SARS-CoV-2 antigens. These preparations may be useful for immediate treatment of COVID-19 disease.

## Introduction

The outbreak of a novel viral respiratory disease, COVID-19, is caused by infection with the Severe Acute Respiratory Syndrome (SARS) Coronavirus 2 (SARS-CoV-2). Due to its extreme transmissibility, COVID-19 has spread dramatically within weeks since the first recognition in China in late December 2019 [1]. Increased human mobility as a global phenomenon has created favorable conditions for COVID-19 to become a pandemic.

Although symptoms are typically mild, in some patient groups COVID-19 can progress to severe respiratory failure which is associated with significant morbidity and mortality. These patients with severe disease are straining the available critical care resources of the most-affected countries [2]. In the short term, the lack of a vaccine and therapeutic agents of proven efficacy against SARS-CoV-2 further aggravates this trend. This critical situation demands a reliable therapy that is immediately available to control the spread of the disease. Convalescent plasma or plasma-derived immunoglobulin (IG), either polyvalent IG (prepared from pooled plasma from thousands of healthy donors) or hyperimmune IG (prepared from the plasma of donors with high titers of antibody against a specific antigen), have been historically used as the fastest therapeutic option in outbreaks of emergent or re-emergent infections [3].

Four main common human coronaviruses have been identified so far: HCoV-229E, HCoV-NL63, HCoV-OC43 and HCoV-HKU1. It is thought that most humans become infected by coronaviruses during their lifetime [4]. SARS-CoV-2 is a novel emerging coronavirus. It joins SARS-CoV, responsible for the SARS outbreak in 2003 and MERS-CoV, responsible for the Middle East respiratory syndrome (MERS) outbreak in 2012. Since coronavirus infections induce virus-neutralizing antibodies, convalescent plasma therapy was successfully used in both SARS [5, 6] and MERS [7] patients.

Common human coronaviruses constantly circulate all around the globe and are accountable for a large proportion of respiratory infections, which in most cases are mild. Because of this ubiquity, antibodies against human common coronaviruses are present in the normal population. Since intravenous IG (IVIG) are polyvalent IG prepared from plasma from thousands of donors, this product covers a large spectrum of immunity of the general population and, as expected, includes anti-coronaviruses antibodies.

It is important to note that coronaviruses of the same subgroup, particularly betacoronaviruses such as HCoV-OC43, HCoV-HKU1, SARS-CoV, SARS-CoV-2, and MERS-CoV, show some cross-reactivity in antigenic responses. Cross-reactivity between SARS-CoV and MERS-CoV with other common human betacoronaviruses has been reported with some neutralization [8–10]. The fact that the new betacoronavirus SARS-CoV-2 is directly related to SARS-CoV (they share more than 90% sequence homology) [11] suggests that antigenic cross-reactivity between them is possible, at least for some proteins.

To explore this potential therapeutic pathway, we designed this study to detect antibodies against common human coronaviruses in IVIG products that may cross-react with the new SARS-CoV-2 virus.

## Material and Methods

### Experimental design

Gamunex^®^-C (Grifols Therapeutics Inc., Raleigh NC, US) and Flebogamma^®^ dual inactivation and filtration (DIF) (Instituto Grifols S.A., Barcelona, Spain) IVIG were tested for cross-reactivity against several betacoronaviruses, including SARS-CoV, MERS-CoV and SARS-CoV-2 antigens, using ELISA techniques.

### IVIG products

Gamunex-C and Flebogamma DIF are highly purified, unmodified human IVIG products manufactured from plasma collected from donors in the US and/or several European countries. Gamunex-C is available in 100 mg/mL (10%) while Flebogamma DIF is available in 50 mg/mL and 100 mg/mL (5% and 10%) IgG concentrations. Both IVIG manufacturing processes contain dedicated steps with high virus clearance capacity, such as solvent/detergent (S/D) treatment, heat treatment, caprylate treatment and Planova™ nanofiltration down to 20 nm pore size.

### Coronaviruses IgG ELISA kits

The following kits were used for the qualitative determination of IgG class antibodies against human coronaviruses: abx052609 Human Coronavirus IgG ELISA kit (Abbexa, Cambridge, UK), against an undetermined antigen; MBS9301037, HCoV-HKU-IgG ELISA kit (MyBioSource, Inc., San Diego, CA, USA), against N protein; DEIA1035. SARS Coronavirus IgG ELISA kit (Creative Diagnostics, Shirley, NY, USA), against virus lysate; RV-402100-1, Human Anti-MERS-NP IgG ELISA Kit (Alpha Diagnostic Intl. Inc., San Antonio, TX, USA), against N protein; RV-402400-1, Human Anti-MERS-RBD IgG ELISA Kit (Alpha Diagnostic Intl. Inc.), against receptor-binding domain (RBD) of S1 subunit spike protein (S1/RBD): RV-402300-1, Human Anti-MERS-S2 IgG ELISA Kit (Alpha Diagnostic Intl. Inc.), against S2 subunit spike protein; RV-405200 (formerly RV-404100-1), Human Anti-SARS-CoV-2 Virus Spike 1 [S1] IgG ELISA Kit (Alpha Diagnostic Intl. Inc.), against S1 subunit spike protein. In all cases the determinations were carried out following the manufacturer’s instructions.

### Sample preparation and testing

IVIG samples were serially diluted using the buffer solutions provided in each IgG ELISA kit. With the IVIG 5% product, the dilution series was: neat (undiluted), 1:5, 1:50, 1:500, 1:1000, and 1:5000. With the IVIG 10% product, the dilution series was: neat, 1:10, 1:100, 1;1000, 1:2000, and 1:10000. Therefore, final IgG concentrations of the samples were: 50 mg/mL, 100 mg/mL, 10 mg/mL, 1 mg/mL, 100 μg/mL, 50 μg/mL, and 10 μg/mL. In SARS-CoV-2 tests, additional dilutions of 1:300 (333 μg/mL) and 1:600 (167 μg/mL) were included.

Reactivity against the coronavirus antigens in the different ELISA kits was rated as negative (−) if no reactivity was observed even with neat IVIG, or positive (+) if the lowest IVIG dilution demonstrated reactivity. The number of test replicates performed was 2-3 for Gamunex-C, 2-4 for Flebogamma 10% DIF, and 1-2 for Flebogamma 5% DIF.

## Results

Both Gamunex-C and Flebogamma DIF showed consistent reactivity to components of the tested viruses including a variety of virus proteins, except for the N-protein from HCoV-HKU1. There was no reactivity to this protein even with undiluted IVIG samples.

As shown in Table 1, positive reactivity was particularly apparent in SARS-CoV, MERS-CoV, and SARS-CoV-2. In the case of MERS-CoV, positive reactivity was observed in IVIG samples down to 1:2000 dilution (50 μg/mL) for N protein, S1-RBD protein and S2 protein. For SARS-CoV-2 S1 protein, positive reactivity ranged from an IVIG concentration of 100 μg/mL with Gamunex-C to 1 mg/mL with Flebogamma 5% DIF (Table 1).

**Table 1.** Results of IgG reactivity against different coronaviruses. Concentration denotes the last IVIG dilution with positive result (+), or no positivity even undiluted (−). N= 1-4 tests.

| IVIG product | Country of origin of the plasma | Virus and antigen/target |  |  |  |  |  |  |  |  |  |  |  |
| --- | --- | --- | --- | --- | --- | --- | --- | --- | --- | --- | --- | --- | --- |
|  |  | HCoV (betacoronavirus) |  | SARS-CoV |  | MERS-CoV |  |  |  |  |  | SARS-CoV-2 |  |
|  |  | Undetermined |  | Culture lysate |  | N protein |  | S1 protein/RBD |  | S protein |  | S1 protein |  |
|  |  | Concentration | Result | Concentration | Result | Concentration | Result | Concentration | Result | Concentration | Result | Concentration | Result |
| Gamunex-C 10% | US | 100 mg/mL | - | 1 mg/mL | + | 100 µg/mL | + | 50 µg/mL | + | 50 µg/mL | + | 100 µg/mL | (+) |
| Flebogamma 5% DIF | US | 50 mg/mL | + | 10 mg/mL | + | 50 µg/mL | + | 50 µg/mL | + | 50 µg/mL | + | 1 mg/mL | (+) |
| Flebogamma 10% DIF | Spain | 100 mg/mL | + | 10 mg/mL | + | 100 µg/mL | + | 50 µg/mL | + | 100 µg/mL | + | 167 µg/mL | (+) |
| Flebogamma 5% DIF | Czech Republic | 50 mg/mL | + | 10mg/mL | + | 1 mg/mL | + | 1 mg/mL | + | 100 µg/mL | + | NT |  |
| Flebogamma 5% DIF | Germany | 100 µg/mL | + | 10 mg/mL | + | 1 mg/mL | + | 1mg/mL | + | 50 µg/mL | + | NT |  |

NT: not tested

Reactivity to HCoV (betacoronavirus undetermined antigen) was also observed, although less consistently: negative for Gamunex, but positive for Flebogamma DIF at low dilutions (Table 1).

## Discussion

The need for readily available effective therapies to combat SARS-CoV-2 infection is compelling. In this study, we considered whether IVIG treatment could contribute to COVID-19 disease management. To test this hypothesis, known currently available IVIG products, Gamunex-C and Flebogamma DIF, were tested for cross-reactivity with SARS-CoV-2 and other coronaviruses, including SARS-CoV and MERS-CoV. We found significant cross-reactivity to components of all tested viruses including the S1 protein of SARS-CoV-2, the protein responsible for virion attachment to the host cell and neutralization [12].

The consistency of our cross-reactivity results among the SARS-CoV-2, SARS-CoV and MERS-CoV viruses, is noteworthy. This replicates with the new SARS-CoV-2, the cross-reactivity already reported for SARS-CoV / MERS-CoV with other human betacoronaviruses [8–10]. Importantly, Gamunex-C and Flebogamma DIF were confirmed to contain antibodies reacting against SARS-CoV-2 antigens, which could be important in the quest for an immediate therapy for COVID-19.

ELISA results for the undetermined antigen of HCoV were also mostly positive. This was in contrast to HCoV-HKU1, which had negative reactivity. HCoV-HKU1 was discovered in 2005 in Hong Kong and, although it did not result in an outbreak and had only restricted spread, this virus is probably still circulating in the population [13, 14].

However, negativity of an IVIG reaction using a single ELISA coronavirus kit does not mean that such IVIG does not contain antibodies against this pathogen. ELISA sensitivity relies on factors such as the antigen used, the sequence, the organism used to produce it, and the amount of material coated. ELISA results should only be compared qualitatively, since comparison of the results between different kits is difficult based on differences in sensitivity, and there is no gold standard for quantification. In addition, there is scarcity of tests for common coronaviruses.

It has been observed that patients that develop a more severe clinical course of SARS-CoV-2 infection have higher plasma levels of proinflammatory cytokines, suggesting a possible cytokine storm associated with the disease severity [15]. IVIG products have been demonstrated to be effective in the treatment of inflammatory disorders [16]. To date, a number of possible mechanisms for the immunomodulatory and anti-inflammatory effects of IVIG therapy have been described [17, 18], including anti-complement effects [19], anti-idiotypic neutralization of pathogenic autoantibodies [20], immune regulation via an inhibitory Fc receptor [16, 21], enhancement of regulatory T cells [22] and inhibition of Th17 differentiation [23]. Thus, IVIG may mediate a wide variety of biological and immunomodulatory effects via various types of blood cells [23]. Altogether, these known immunomodulatory effects of IVIG products could be beneficial in COVID-19 disease management. Anecdotally, there was a case series report describing a positive effect of high dose IVIG in three patients with COVID-19 [24]. It should be noted that IVIG products are generally deemed safe and well-tolerated. Most of their adverse effects are mild and transient [25].

In conclusion, we consistently observed cross-reactivity of IVIG products with SARS-CoV-2, SARS-CoV and MERS-CoV using ELISAs from different manufactures. This evidence supports the presence of anti-SARS-CoV-2 cross-reacting antibodies in these IVIG preparations. These results together with the known immune properties of IVIG, suggest a potential positive contribution of currently available IVIG products to COVID-19 disease management. Further steps to confirm the functionality of IVIG antibodies such are neutralization studies are warranted.

## Acknowledgements

Jordi Bozzo PhD, CMPP (Grifols) is acknowledged for medical writing and Michael K. James PhD (Grifols) is acknowledged for editorial support in the preparation of this manuscript. Contribution from Antonio Páez MD, Sandra Fernández MD, and Elisabeth Calderón PhD (Grifols) who provided their expert opinion is acknowledged. The authors acknowledge the expert technical assistance from Daniel Casals, Eduard Sala, Judith Luque and Gonzalo Mercado (Grifols, Viral and Cell Culture Laboratory)

